## Supplementary Materials for "The genomic architecture of blood metabolites based on a decade of genome-wide analyses"

### Supplementary Notes

#### Supplementary Note 1. Overview of GWA and (exome-) sequencing studies

We updated and extended the existing review for QTL discovery^1^ for metabolites in European cohorts and supply the complete list of references below:

| **Study** | **Biofluid** | **Sample (N)** | **Population** | **Study type** | **Genome build** | **Metabolomics platform(s)** | **Metabolites (N)** |
| --- | --- | --- | --- | --- | --- | --- | --- |
| (Gieger et al. 2008)^2^ | Serum | 284 | European | GWA | NCBI build 35 | Biocrates (MS) | 363 + ratios |
| (Tanaka et al. 2009)^3^ | Plasma | 1,210 + 1,076 | European | GWA | NCBI build 35 | gas chromatography | 6 |
| (Hicks et al. 2009)^4^ | Plasma & serum | 4,400 | European | GWA | NCBI build 36 | Lipidomics (MS) | 33 + 43 ratios |
| (Illig et al. 2010)^5^ | Serum | 1,809+422 | European | GWA | NCBI build 36 | Biocrates (MS) | 163 + 26,406 ratios |
| (Lemaitre et al. 2011)^6^ | Plasma | 8,866 | European | GWA | NCBI build 35 | gas chromatography | 4 |
| (Suhre, Shin, et al. 2011)^7^ | Serum | 1,768 + 1,052 | European | GWA | NCBI build 36 | Metabolon (MS) | 276 + 37,179 ratios |
| (Nicholson et al. 2011)^8^ | Urine & plasma | 211 | European | GWA | NCBI build 37 | NMR in urine & Biocrates (MS) in plasma | 512 urine + 163 + ratios plasma |
| (Kettunen et al. 2012)^9^ | Serum | 8,330 | European | GWA | NCBI build 36 | Nightingale Health (NMR) | 117 + 99 ratios |
| (Demirkan et al. 2012)^10^ | Plasma | 4,043 | European | GWA | NCBI build 36 | Lipidomics (MS) | 153 |
| (Tukiainen et al. 2012)^11^ | Serum | 8,330 | European | refinement known loci | NCBI build 36 | Nightingale Health (NMR) | 117 + 99 ratios |
| (Krumsiek et al. 2012)^12^ | Serum | 1,768 | European | GWA | NCBI build 36 | Metabolon (MS) | 517 |
| (Wu et al. 2013)^13^ | Plasma | 8,961 | European | GWA | NCBI build 36 | gas chromatography | 4 |
| (Raffler et al. 2013)^14^ | Plasma | 1,757 | European | GWA | NCBI build 36 | NMR + Biocrates + Metabolon (MS) | 8,600 + 124,750 ratios |
| (Hong et al. 2013)^15^ | Serum | 402 + 489 | European | GWA | NCBI build 36 | MS | 6,138 |
| (Xie et al. 2013)^16^ | Plasma | 1,004 + 341 | European | GWA | NCBI build 36 | Metabolon (MS) | 14 |
| (Rhee et al. 2013)^17^ | Plasma | 2,076 | European | GWA | NCBI build 36 | MS | 217 |
| (Shin et al. 2014)^18^ | Serum | 7,824 | European | GWA | NCBI build 36 | Metabolon (MS) | 486 + 98,346 ratios |
| (Guan et al. 2014)^19^ | Plasma | 8,631 | European | GWA | NCBI build 36 | gas chromatography | 5 |
| (Mozaffarian et al. 2015)^20^ | Red blood cells + plasma | 8,013 | European + African American + Chinese + Hispanic | GWA | NCBI build 36 | gas chromatography | 5 |
| (Lemaitre et al. 2015)^21^ | Plasma | 10,129 | European | GWA | NCBI build 36 | gas chromatography | 3 |
| (Demirkan et al. 2015)^22^ | Serum | 2,118 | European | GWA + exome sequencing | NCBI build 36 (GWA) and 37 (exome) | ‘Leiden’ (NMR) | 42 |
| (Yu et al. 2015)^23^ | Serum | 1,152 + 718 + 753 | African American + European | exome sequencing + exome chip | NCBI build 37 | Metabolon (MS) | 1 |
| (Draisma et al. 2015)^24^ | Serum | 7,478 + 1,182 | European | GWA | NCBI build 36 | Biocrates (MS) | 129 |
| (Tintle et al. 2015)^25^ | Red blood cells | 2,633 | European | GWA | NCBI build 36 | gas chromatography | 14 |
| (Hu et al. 2016)^26^ | Red blood cells + plasma | 2,685 + 656 + 8,866 + 8,962 | Chinese + European | GWA | NCBI build 36 | gas chromatography | 9 |
| (Hartiala et al. 2016)^27^ | Plasma | 1,985 + 1,895 + 400 | European | GWA | NCBI build 36 | MS | 9 |
| (Kettunen et al. 2016)^28^ | Serum & plasma | 24,925 | European | GWA | NCBI build 37 | Nightingale Health (NMR) + ‘Leiden’ (NMR) | 123 |
| (Yet et al. 2016)^29^ | Serum | 1,001 | European | GWA | NCBI build 36 | Biocrates & Metabolon (MS) | 648 |
| (Yazdani et al. 2016)^30^ | Serum | 1,456 | European | whole genome sequencing | NCBI build 37 | Metabolon (MS) | 16 |
| (Fall et al. 2016)^31^ | Plasma | 1,138 + 970 + 1,630 | European | GWA + look-up | only rsids used | non-targeted MS | 15 |
| (Rhee et al. 2016)^32^ | Plasma | 2,076 + 1,528 | European | exome chip | NCBI build 37 | Metabolon (MS) + other MS | 217 |
| (Yu et al. 2016)^33^ | Serum | 1,872 + 1,552 | African American + European | exome + whole genome sequencing | NCBI build 37 | Metabolon (MS) | 70 |
| (Lotta et al. 2016)^34^ | Plasma | 16,596 | European | GWA | NCBI build 37 | Biocrates (MS) + Metabolon (MS) | 3 |
| (Hu et al. 2017)^35^ | Red blood cells + plasma | 3,521 + 12,020 | Chinese + European | GWA | NCBI build 36 | gas chromatography + GC-MS | 6 |
| (Long et al. 2017)^36^ | Serum | 1,960 | European | whole genome sequencing | NCBI build 38 | Metabolon (MS) | 644 |
| (Davis et al. 2017)^37^ | Serum | 8,372 | European | exome chip | NCBI build 37 | Nightingale Health (NMR) | 72 |
| (Teslovich et al. 2018)^38^ | Serum | 8,545 + 2,591 | European | GWA + exome chip | NCBI build 37 | Nightingale Health (MS) | 9 + 36 ratios |
| (Feofanova et al. 2018)^39^ | Serum | 1,872 + 1,552 | European + African American | whole exome + whole genome sequencing | NCBI build 37 | Metabolon (MS) | 102 |
| (Kalsbeek et al. 2018)^40^ | Red blood cells | 2,374 | European | GWA | NCBI build 37 | gas chromatography | 22 + 15 ratios |
| (de Oliveira Otto et al. 2018)^41^ | Red blood cells + plasma | 11,494 | European | GWA | only rsids used | gas chromatography + GC-MS | 5 |

#### Supplementary Note 2. four-variance component models with HMDB classes

Metabolite SNPs were curated from 40 GWA and (exome-) sequencing studies (**Supplementary Note 1**; **Supplementary Data 1**) and all metabolites were categorized by HMDB ‘super class’, ‘class’ and ‘subclass’ (see **Methods**). For each class represented in our metabolite data (12) we created GRMs capturing solely the metabolite loci for metabolites of this specific class (*h^2^_Class-hits_*). Metabolite loci for each class were identified by including all published metabolite-SNP associations of the relevant class in a clumping procedure (r^2^ = 0.1, radius = 500kb). The lead SNPs identified by the clumping procedure were then used as input to create LD-corrected GRMs in LDAK (version 4.9). We used the same technique to create corresponding ‘non-class’ GRMs, which used included the lead SNPs of all metabolite-SNP associations, excluding the relevant class loci and LD proxies (*h^2^_notlass-hits_*). This resulted in 4-variance component models, as resolved in the GCTA software (version 1.91.7). **Figure 1** provides an overview of the four-variance component models, including the specification of the underlying GRMs.

The 4-variance component models for the 12 different classes of metabolites had high degrees of non-convergence (37.9% total; **Supplementary Table 2**). Non-convergence of the 4-variance component models appears to be associated with the number of SNPs in the ‘class-specific’ GRMs, with GRMs with a small number of SNPs displaying more convergence issues (**Supplementary Table 2**). To address the convergence issues we recoded the HMDB metabolite classes into HMDB metabolite super classes, reducing the 12 classes to 5 classes (**Supplementary Table 4**). Metabolites belonging to the organic nitrogen compounds, organic oxygen compounds and protein classes were not analyzed as these super classes consisted of only a single class, which had poor convergence. Combining classes into 2 super classes with a larger number of SNPs in the GRM results in a convergence rate of 97.8% (**Supplementary Table 4**). The complete results for the final 4-variance component models have been described in the main text of the manuscript.

#### Supplementary Note 3. Comparison of weighted vs. unweighted GRMs and clumped vs. not-clumped GRMs

##### Simulations of using a weighted vs. unweighted GRMs

Genome-wide complex trait analysis (GCTA)^46^ was used to simulate 20 traits with heritabilities of 20%, 40%, 60% and 80% under the assumption that 1000 random single nucleotide polymorphisms (SNPs; (e.g., polygenic traits) or 5 random SNPs (e.g., near monogenic traits) are causally underlying these traits. Data underlying these simulations were the cross-imputed genotype sets (see **Methods**). SNP heritability (h^2^_SNP_) analyses were run in GCTA for all simulated traits using a weighted genetic relatedness matrix (GRM; LDAK^43,44^) and an unweighted GRM in order to compare the accuracy of the h^2^_SNP_ estimates. The simulations show that estimated heritabilities from both GRMs deviate slightly from the simulated heritability.

Under the polygenic assumption, the weighted GRM shows an upwards bias for lower simulated heritabilities (**Supplementary Figure 4a**), while the unweighted GRM shows a larger downwards bias for higher simulated heritabilities (**Supplementary Figure 4b**). Under near monogenic assumptions the differences between the weighted and unweighted GRMs largely disappear (**Supplementary Figure 4c-4d**). The accuracy for both GRMs under near monogenic assumptions appeared to be quite similar, therefore, additional simulations under near monogenic assumptions were run as previous studies have shown that relatively large proportions of variance in serum metabolite levels can be explained by a relatively small number of genetic variants^2,5,9,24^. Three sets of 5 ‘causal’ SNPs were selected, with one set of SNPs in low linkage disequilibrium (LD) with each other, a set of medium LD SNPs and a set of SNPs in high LD. High, medium and low LD was assigned based on the LD-weighting as calculated by LDAK. For each of these SNP sets 20 traits with heritabilities of 10-50% were simulated and h^2^_SNP_ estimates were compared for the weighted and unweighted GRMs. Across all LD sets the unweighted GRM underestimated the simulated heritability, weighted GRMs appeared to have smaller biases (**Supplementary Figure 5**). However, as genotyped or imputed datasets as used here already includes the LD-structure in the data and likely biases the simulations in favour of weighted GRMs, we are only able to conclude that, as our simulations are likely biased to favor LDAK, LDAK does at least as well as GCTA.

##### Clumped vs. not-clumped weighed and unweighted GRMs

After curating all metabolite-SNP associations of the past decade we have identified 35,138 unique metabolites SNPs (see **Methods** and **Supplementary Figure 1**). Many of these SNPs are from the same genetic loci and likely describe the same association for different metabolites. Including all SNPs may induce upward bias of the heritability estimates. Therefore, in constructing the weighted GRMs from the class-specific and not-class-specific metabolite loci we first clump all metabolite-SNP associations (**Supplementary Figure 1**) to obtain independent metabolite loci. However, clumping of SNPs before creating weighted GRMs may potentially remove real signal and result in underestimation of the heritability estimates. To investigate if clumped indeed results in downward bias of our heritability estimates we compared clumped and not-clumped LDAK metabolite GRMs. We used the GCTA power calculator to compare the estimated SE’s and power for the different GRMs included in our manuscript using clumped and not-clumped SNPs (<https://cnsgenomics.shinyapps.io/gctaPower/>).

The differences between the GRMs using clumped or not-clumped SNPs are very small (**Supplementary Table 16**). Across all 5 GRMs the average mean differences between the clumped and not-clumped GRMs are negative, indicating that clumped GRMs results in smaller SE’s, though with a minimal difference (**Supplementary Table 16**). Similarly, the average power differences are zero or positive (again with minimal differences), indicating that clumped GRMs are as powerful or more powerful than not-clumped GRMs (**Supplementary Table 16**). While the gain of using a clumped weighted LDAK GRM versus a not-clumped LDAK GRM is small, we feel that the given the large SE’s we’ve observed in our study, the choice for clumped weighted LDAK GRMs is valid. However, by clumping the SNPs prior to creating the weighed GRM the differences between weighted and unweighted GRMs become very small (**Supplementary Table 17**). A final worry about the use of a weighted GRM is the effect of thresholding^45^ in order to obtain the GM including only the closely related individuals. As can be seen in the scatterplot (**Supplementary Figure 6**) of the weighted (LDAK) vs unweighted (GCTA) thresholded GRM, there are some discrepancies in the individuals identified as not closely-related (set to zero) in one or the other methods (horizontal/vertical outliers bottom left corner graph). However, the number of relationships not classified as unrelated by both methods is very low. Fourteen pairs were classified as unrelated by the LDAK method, but not by GCTA, and two pairs were classified as unrelated by the GCTA method, but not by LDAK. Therefore, the effect of using LDAK or GCTA to create the GRM retaining only the relationships among closely related individuals will be minimal.

#### Supplementary Note 4. Covariate determination

##### Generalized estimation equation models

The contribution of covariates to the metabolite levels after quality control was estimated using generalized estimation equation models (GEE). GEE models were fitted in the R programming language (R version 3.5.1) using the GEE-package (version 4.13-19) with the Gaussian link function for continuous data and the ‘exchangeable’ option to correct for the correlation structure due to family resemblance with 100 iterations. All GEE models were done on a subset of the data in which individuals with missing data on the covariates were excluded. Matrix Spectral Decomposition (MSD)^48^ was applied to estimate the number of independent variables in the correlation matrix of all metabolites that passed quality control (QC). Correction for multiple testing was done by a Bonferroni correction for the number of independent variables (α= 0.05/ N independent variables), as in van Dongen et al. (2015)^49^. MSD identified 93 independent variables, therefore, the significance threshold was *p* ≤ 0.0005.

Covariates included sex, age at blood draw, body mass index (BMI), smoking status (never, former, current), use of sex-hormones (y/n, defined as medications belonging to the ‘G03 – sex hormones and modulators of the genital system’ ATC class), use of systemic hormones (y/n, here systemic hormone use is defined as including medication belong to the following ATC classes: ‘H01 – pituitary and hypothalamic hormones and analogues’, ‘H02 – corticosteroids for systemic use’, ‘H03 – thyroid therapy’, and ‘H04 – pancreatic hormones’), use of medication for diabetes (‘A10 – drugs used in diabetes), use of medication for hypertension (y/n, defined as medications belong to the following ATC classes: ‘C02 – antihypertensives’. ‘C03 – diuretics’, ‘C04 – peripheral vasodilators’, ‘C05 – vasoprotectives’, ‘C07 – beta blocking agents’, ‘C08 – calcium channel blocker’, and ‘C09 – agents acting on the renin-angiotensin system’), use of medication for chronic obstructive pulmonary disease (COPD; y/n, defined as medications belong to the ATCH classes ‘R01 – nasal preparations’ and ‘R03 – drugs for obstructive airway diseases), and use of anti-inflammatory medications (y/n, defined as medications belonging to the following ATC classes: ‘L01 – antineoplastic agents’, ‘L02 – endocrine therapy’, ‘L03 – immunostimulants’, ‘L04 – immunosuppressants’, ‘M01 – anti-inflammatory and antirheumatic products’ and ‘M02 – topical products for joint and muscular pain’). We also considered the following technical covariates: genotyping chip (‘AXIOM’, ‘AFFY6’, ‘ILL660’, ‘ILL1M’, ‘ILLGSA’, ‘PERAFF’ or ‘GONL sequencing’), measurement batch (not included for the UPLC-MS Lipidomics platform) and population stratification by including the first 10 genetic principal components (PC) as based on the Dutch population^50^.

Metabolite levels showed associations with sex, age, BMI, smoking status and use of sex-hormones. After correction for multiple testing 79.9% of the metabolites had a significant association with age (295/369), 78.6% with sex (290/369), 50.4% with use of sex-hormones (186/369), 48% with BMI (177/369) and 32.8% with smoking status (121/369; **Supplementary Table 18**). For the significantly associated metabolites, we observed overall higher metabolite levels for older individuals, individuals with a higher BMI and individuals using sex-hormones (**Supplementary Table 18**). Overall lower metabolite levels were observed for current smokers as compared with former or never smokers, while the effect of sex on metabolite levels varied greatly (**Supplementary Table 18**). Only few metabolites were associated with diabetes medication (0.8%), anti-hypertension medication (3.3%) or the use of systemic hormones (1.6%; **Supplementary Table 18**). COPD medication and anti-inflammatory medication were not significantly associated with metabolite levels of any of the four platforms after correction for multiple testing (**Supplementary Table 18**).

##### Comparing three different two-variance component models in GCTA

In order to determine the influence of covariates on the heritability estimates we ran three different models. Aschard et al. (2015) showed that including a heritable trait as covariate in GWAS analysis, or when calculating heritability for a trait, might lead to biased estimates for the trait of interest^51^. For this reason we compared the ‘full’ and ‘reduced’ two-variance component models, both included (potentially) heritable covariates, with a ‘sparse’ two-variance component model, that did not include any (potentially) heritable covariates. The ‘full’ model included all covariates as were included in the GEE models, the ‘reduced’ model excluded COPD medication use, anti-inflammatory medication use and use of systemic hormones, as these were overall not significantly associated with metabolite levels across the different platforms, and finally the ‘sparse’ model included only sex and age at blood draw. In addition all models also included genetic PCs, genotyping chip and when applicable measurement batch (see **Supplementary Table 12**).

The total heritability (*h^2^_total_*) and *h^2^_SNP_* estimates for all metabolites across all models can be found in **Supplementary Table 19**. The mean and median *h^2^_total_* and *h^2^_SNP_* estimates across all metabolites per model are provided in **Supplementary Table 13**. Aschard et al. (2015) predicted that models including heritable covariates will have increased heritability estimates as compared with the models without heritable covariates. In contrast to this prediction, we observed slightly higher mean and median *h^2^_total_* and *h^2^_SNP_* estimates when considering the ‘sparse’ model as compared with the ‘full’ and/or ‘reduced’ model. These results seem to imply that the inclusion of (potentially) heritable traits generally does not bias the estimates in an upwards fashion for the metabolomics platforms under consideration.

The Log Likelihood (LogL) for each of the models for all metabolites across all four platforms have been extracted to calculate the likelihood ratio test (LRT) to compare the full model with the reduced and sparse models (**Supplementary Table 19**). The LRT shows that on average (99.2%) the reduced model is not a better fit for the data as compared to the full model, but the sparse model generally fits the data better (56.9%; **Supplementary Table 19**). Therefore, we decided to use the most sparse model in the further analyses.

### Supplementary Figures

**Supplementary Figure 1.** Flowchart describing the filtering of metabolite SNPs and GRM construction for the 4-variance component models.

This flowchart describes how the 242,580 metabolite-SNP associations as identified from GWA and rare-variant analyses (**Supplementary Note 1**; **Supplementary Data 1**) were converted to NCBI build 37, extracted for NTR participants from the 1000GP3 imputed data and filtered on MAF, HWE and R^2^ (blue boxes at top of the figure indicated by the red curly bracket). The metabolite-SNP associations of the filtered SNPs were clumped (r^2^ = 0.10) to obtain the metabolite loci and LD-proxies of the lipid and the organic acids, respectively (blue). To obtain the non-superclass loci, the superclass-specific loci and LD-proxies were removed from the overall list of metabolite-SNP associations and prior to clumping (blue). The lipid-loci, not-lipid loci, organic acid loci and not-organic acid loci give rise to four GRMs, respectively, as indicated by the black boxes and arrows in the flowchart. The two additional GRMs included in the 4-variance component GREML models are based on the cross-platform imputed SNPs (see **Methods**), where the lipid and organic acid loci, LD-proxies and 50 kb surrounding these SNPs have been removed from one of the cross-platform GRMs (black box in flowchart).

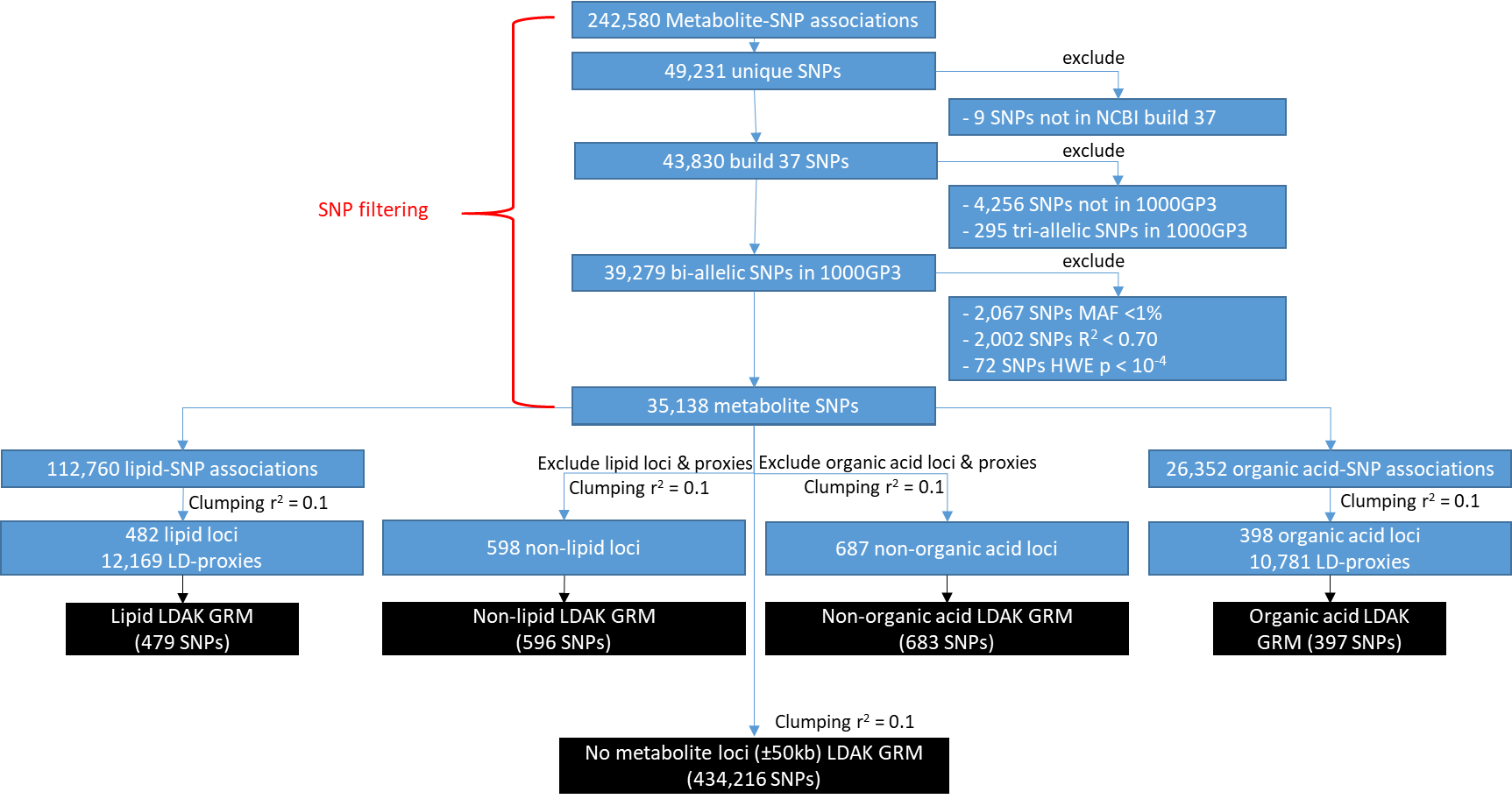

**Supplementary Figure 2.** Heritability estimates of *h^2^_total_*, *h^2^_Class-hits_*, *h^2^_Notclass-hits_* and h^2^_Metabolite-hits_ with standard errors for the phosphatidylcholines (PCs) and triglycerides (TGs) ordered by the number of carbon atoms and double bonds in each species. (**a**) PCs as measured on the Biocrates platform. (**b**) PCs as measured on the UPLC-MS lipidomics platform. (**c**) TGS as measured on the UPLC-MS lipidomics platform. **Supplementary Table 3** provides the estimates for each of the individual metabolites.

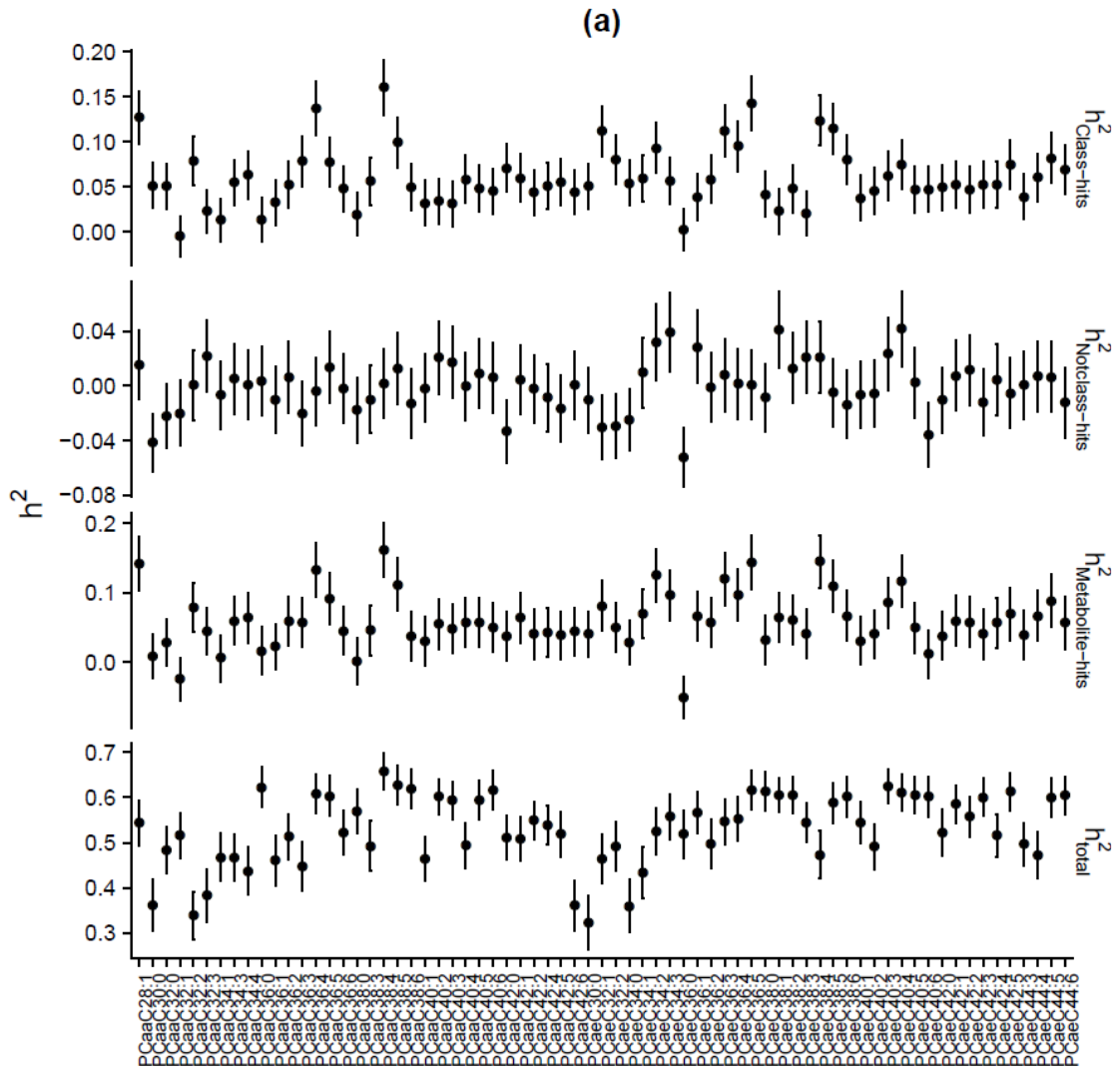

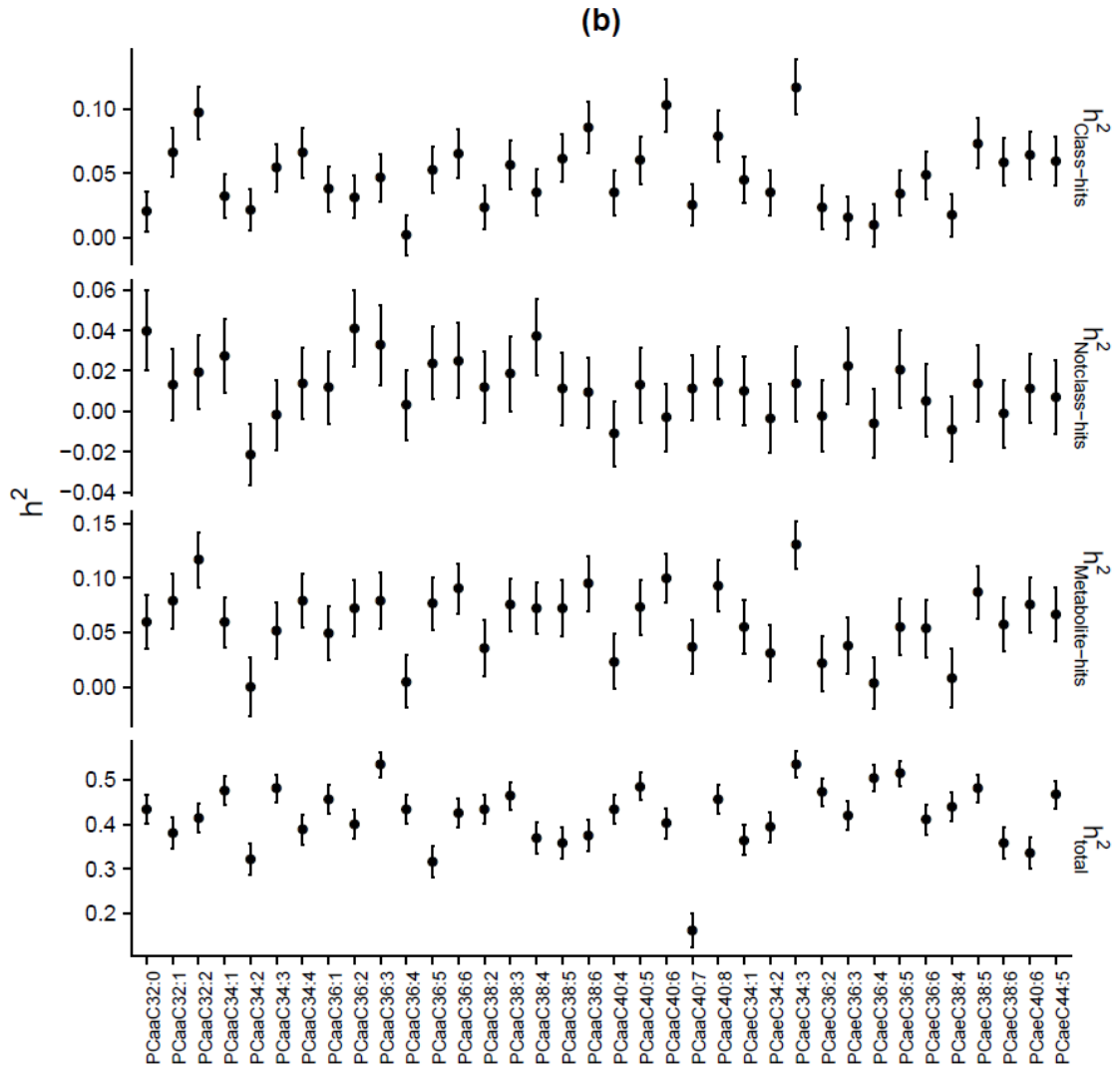

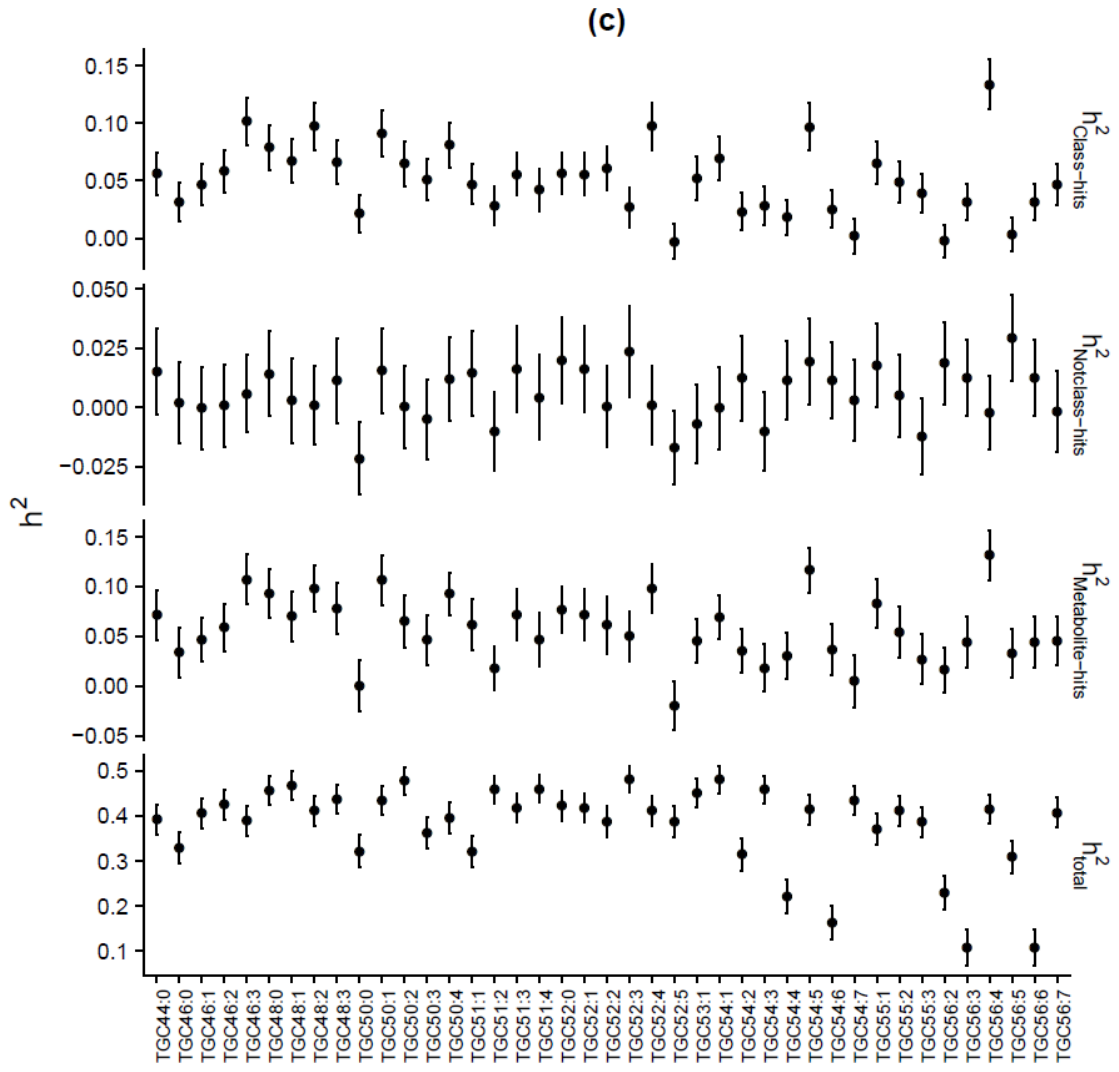

**Supplementary Figure 3.** Histogram of the off-diagonal elements of the Genetic Relatedness Matrix (GRM) of all participants. The off-diagonal elements represent the pairwise relatedness for all participants. **(a)** Histogram of all off-diagonal elements of the LDAK GRM for all participants. As a large proportion of relationships among the participants is unrelated (<0.05) this histogram is highly skewed to the left, making the relationships among related individuals undistinguishable. **(b)** Histogram of the off-diagonal elements in the LDAK GRM for the unrelated (<0.05) participants. **(c)** Histogram of all off-diagonal elements of the LDAK GRM for all related (>0.05) participants.

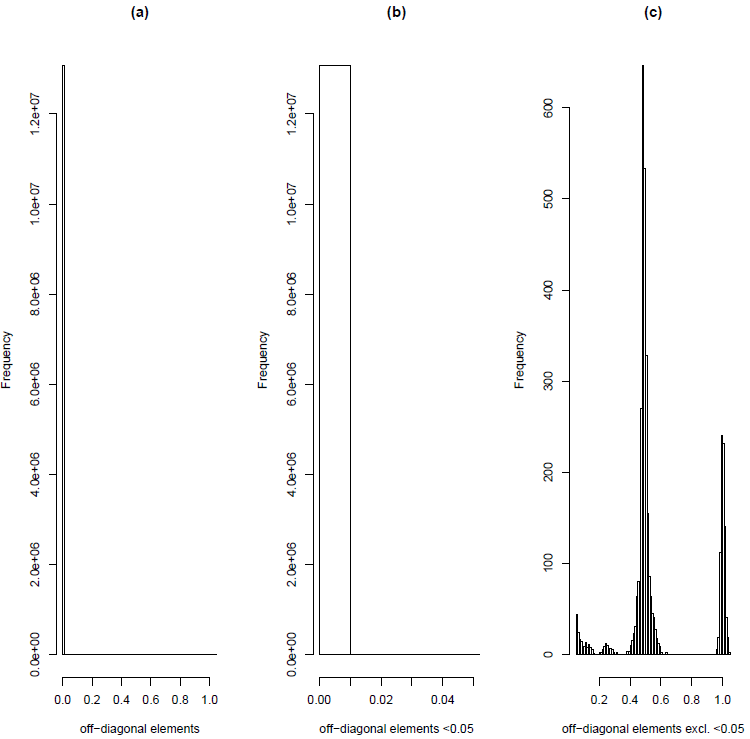

**Supplementary Figure 4.** GCTA results with weighted versus unweighted GRMs of 20 simulated traits under multiple heritabilities with 1000 causal SNPs or 5 causal SNPs. **(a)** GCTA results with unweighted GRM of 20 simulated traits under multiple heritabilities with 1000 causal SNPs. **(b)** GCTA results with weighted GRM of 20 simulated traits under multiple heritabilityies with 1000 causal SNPs. **(c)** GCTA results with unweighted GRM of 20 simulated traits under multiple heritabilities with 5 causal SNPs. **(d)** GCTA results with weighted GRM of 20 simulated traits under multiple heritabilities with 5 causal SNPS. The red line indicates a perfect overlap between the simulated and estimated heritabilities.

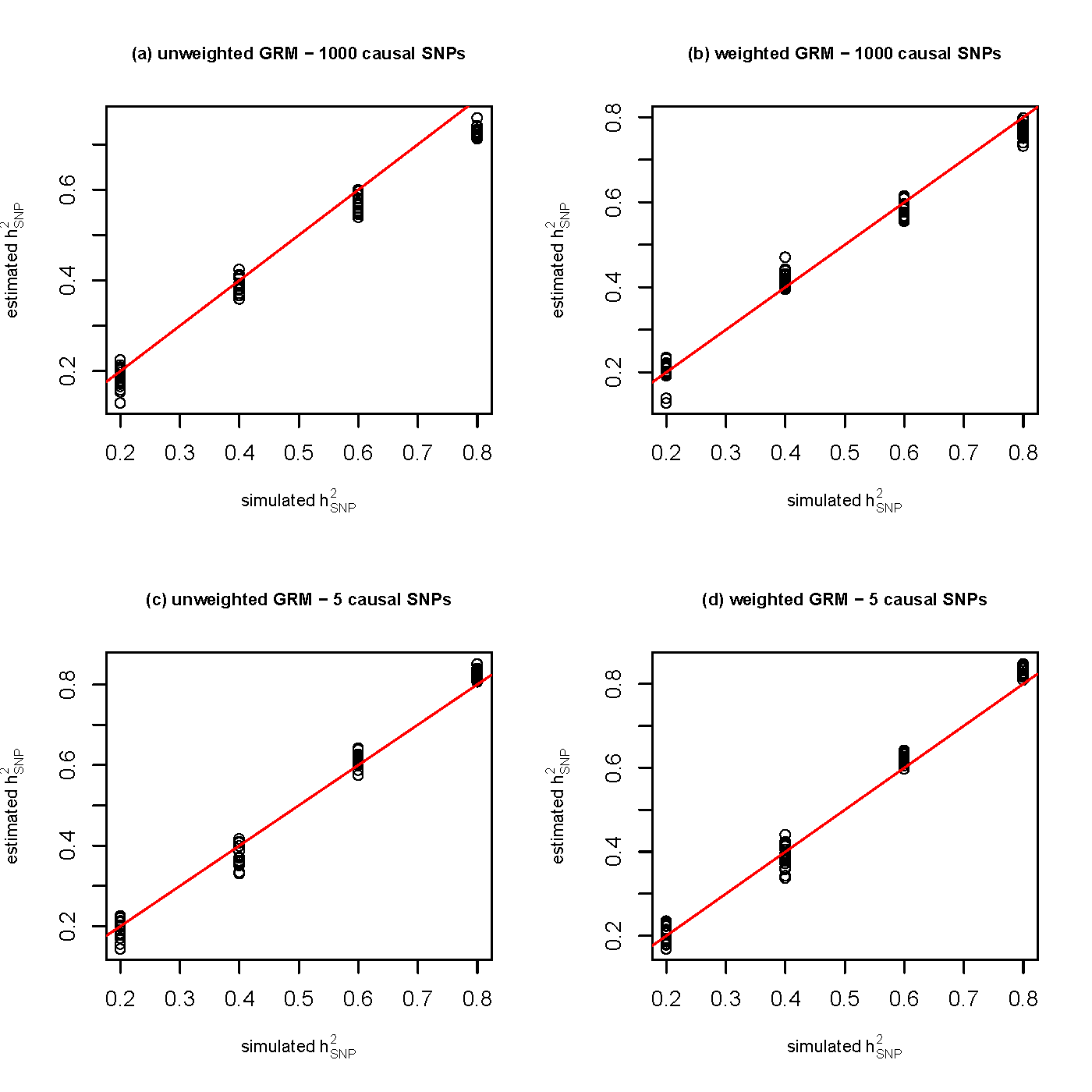

**Supplementary Figure 5.** GCTA results with weighted versus unweighted GRMs of 20 simulated traits under multiple heritabilities with 5 causal SNPs selected to be in low, medium or high LD**.** **(a)** GCTA results with unweighted GRMs of 20 simulated traits under multiple heritabilities with 5 causal SNPs selected to be in low LD. **(b)** GCTA results with weighted GRMs of 20 simulated traits under multiple heritabilities with 5 causal SNPs selected to be in low LD. **(c)** GCTA results with unweighted GRMs of 20 simulated traits under multiple heritabilities with 5 causal SNPs selected to be in medium LD. **(d)** GCTA results with weighted GRMs of 20 simulated traits under multiple heritabilities with 5 causal SNPs selected to be in medium LD. **(e)** GCTA results with unweighted GRMs of 20 simulated traits under multiple heritabilities with 5 causal SNPs selected to be in high LD. **(f)** GCTA results with weighted GRMs of 20 simulated traits under multiple heritabilities with 5 causal SNPs selected to be in high LD. The red line indicates a perfect overlap between the simulated and estimated heritabilities.

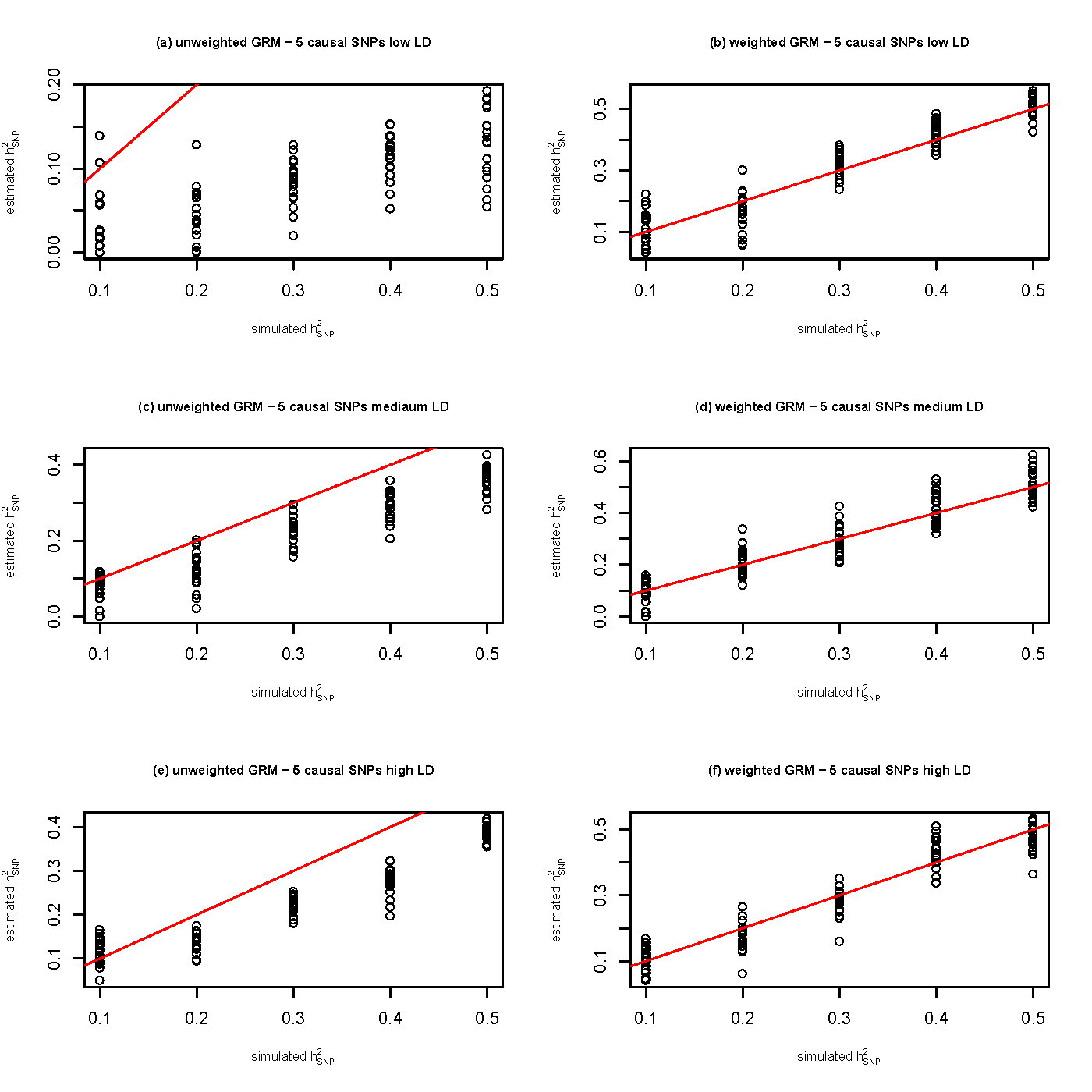

**Supplementary Figure 6.** Scatterplot of the off-diagonal elements of the clumped LDAK versus clumped GCTA GRMs with the relationships for unrelated individuals (<0.05) set to zero. **(a)** Scatterplot of all off-diagonal elements of the clumped LDAK versus clumped GCTA GRMs with the relationships for unrelated individuals (<0.05) set to zero. **(b)** **.** Scatterplot of the <0.05 off-diagonal elements of the clumped LDAK versus clumped GCTA GRMs with the relationships for unrelated individuals (<0.05) set to zero.

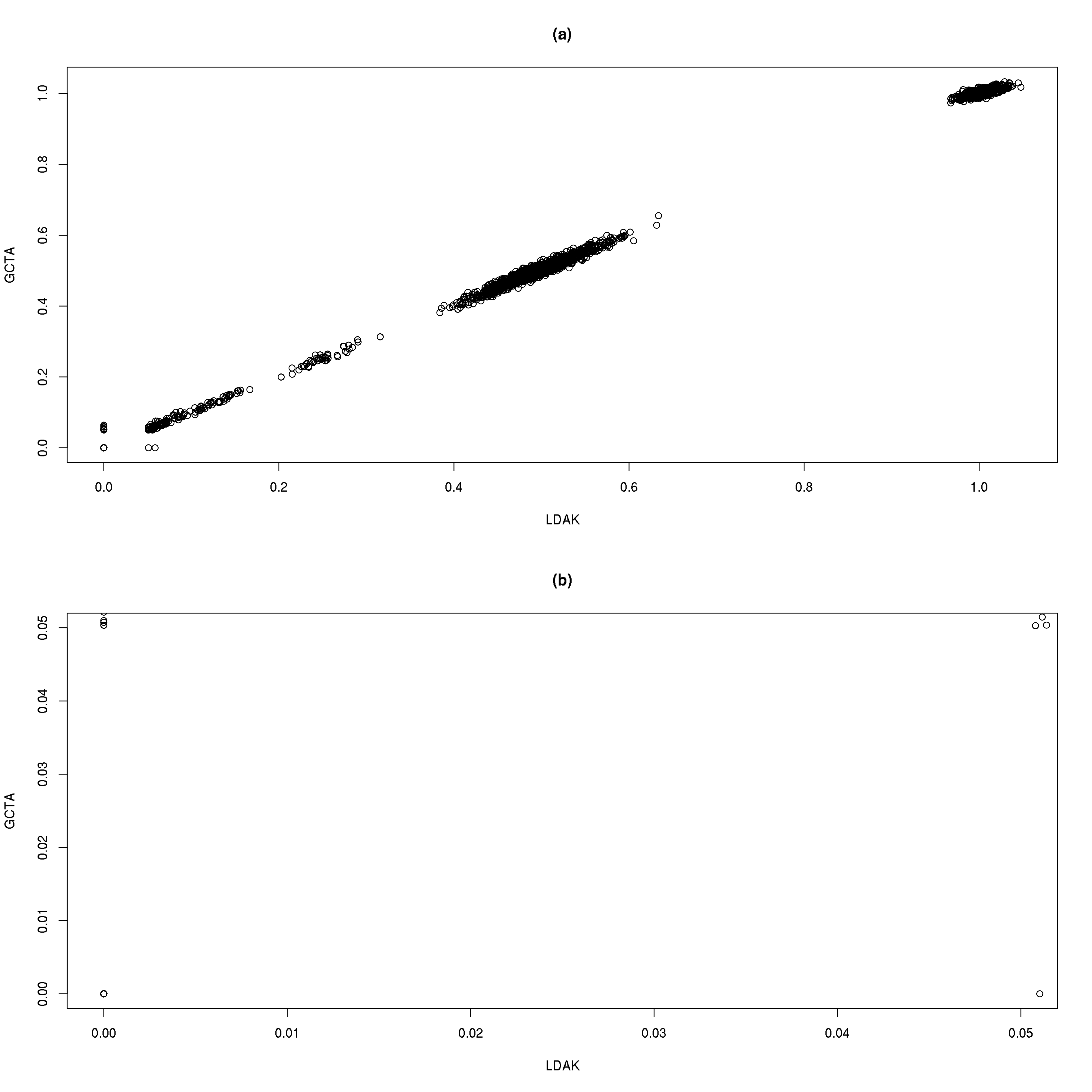
